# Biomarkers in cerebrospinal fluid sediments

**DOI:** 10.1101/2025.06.10.657984

**Authors:** Raquel Alsina, Marta Riba, Marina Sartorio, Clara Romera, Berta Vilaplana, Eva Prats, Laura Molina-Porcel, Jaume del Valle, Carme Pelegrí, Jordi Vilaplana

## Abstract

Cerebrospinal fluid (CSF) biomarkers for neurodegenerative diseases have been extensively studied over the years. However, CSF samples are routinely centrifuged, and the resulting sediment or pellet is typically discarded to remove cellular debris and high-density particles. This standard practice raises a critical question: could these discarded sediments harbour potential biomarkers relevant to the diagnosis and prognosis of certain brain diseases? In this study, we analysed CSF pellets from various cases and identified, entrapped among undetermined remnants, brain-derived structures such as wasteosomes and psammoma bodies. Furthermore, we observed that disease-relevant proteins can become deposited in the sediment, as is the case for both tau and Aβ_42_ in Alzheimer’s disease or tau in progressive supranuclear palsy disease. These findings suggest that some potential biomarkers might accumulate or be hidden in the sediment and, taken as a whole, the results underscore the need to broaden the scope of biomarker research.

Graphical abstract

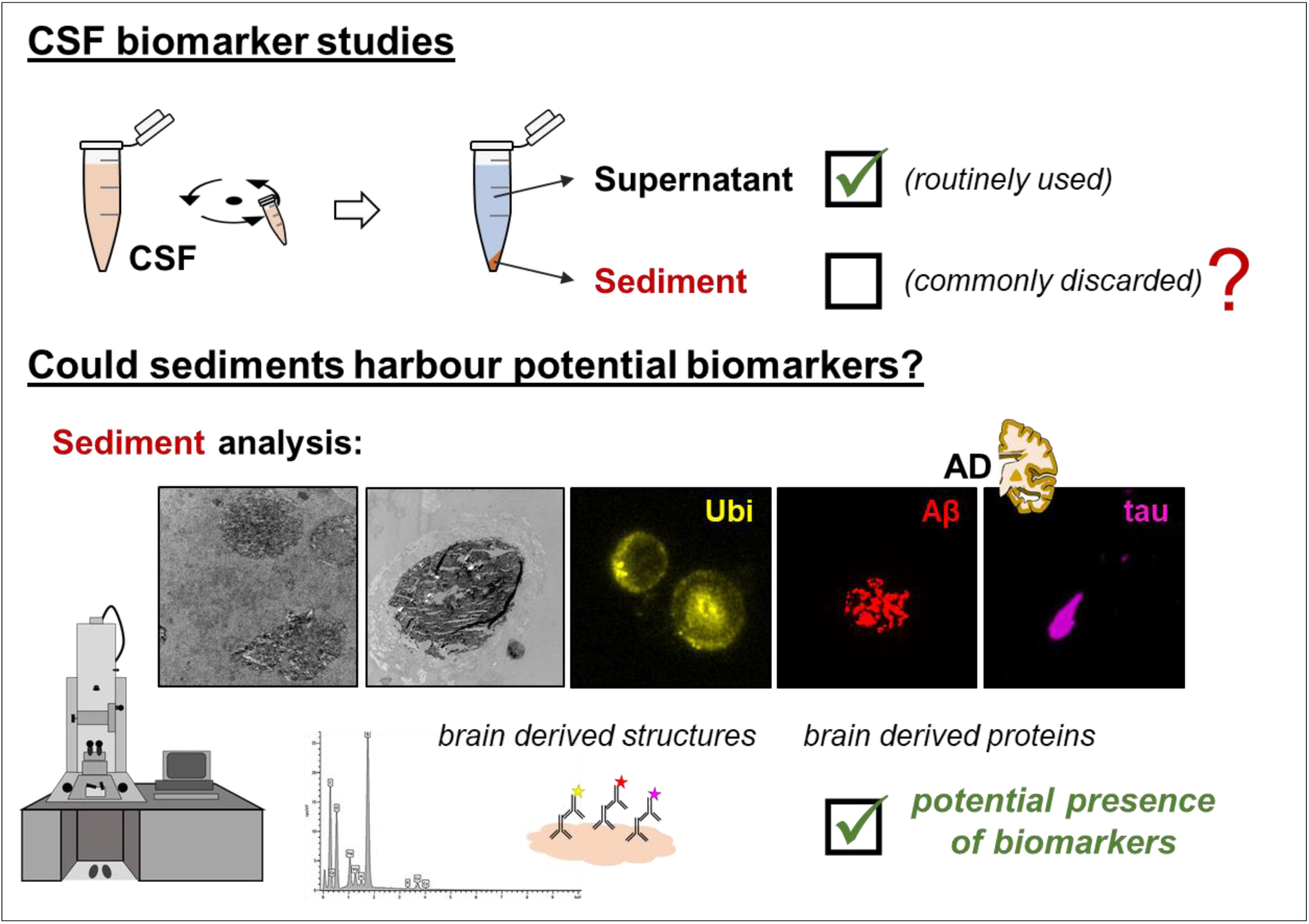

## Introduction

Neurodegenerative diseases are a heterogeneous group of neurological disorders characterized by the progressive loss of neurons in the central or peripheral nervous system. The loss of neurons and the consequent collapse of the structure and function of neural networks result in the breakdown of the core communicative circuitry, culminating in impaired memory, cognition, behaviour, sensory, and/or motor function [1].

Among these diseases, Alzheimer’s disease (AD) stands as the most prevalent, followed by Parkinson’s disease and frontotemporal lobar degeneration (FTLD). All these diseases share a common hallmark as they entail the formation of insoluble aggregates of specific proteins which are often implicated in the pathology and progression of the diseases [2,3]. In the case of AD, amyloid-β (Aβ) and tau aggregates constitute the major components of Aβ plaques and neurofibrillary tangles respectively [4]. In Parkinson’s disease, aggregates of α-synuclein are the major component of Lewy bodies present in the brains of these patients [5]. Finally, for FTLD, aggregates of hyperphosphorylated tau, transactive response DNA-binding protein 43 (TDP-43), and inclusions of fused in sarcoma (FUS) protein have been reported, corresponding to the subtypes FTLD-tau, FTLD-TDP, and FTLD-FUS, respectively [6,7].

Altogether, neurodegenerative diseases are affecting the lives of millions of people worldwide, and their prevalence is expected to rise with increasing life expectancy in most countries [8]. Nonetheless, despite the increasing demand for curative treatments, current therapies remain primarily symptomatic. An exception is found in AD, for which the U.S. Food and Drug Administration (FDA) and the European Medicines Agency have recently approved a drug based on an anti-amyloid monoclonal antibody, Lecanemab (Leqembi®), for use in early-stage AD patients. Lecanemab demonstrated a significant slowing of cognitive decline in early AD in clinical trials and became the first disease-modifying drug to receive traditional FDA approval [9,10]. Additionally, two more drugs based on anti-amyloid monoclonal antibodies, Aducanumab (Aduhelm®) and Donanemab (Kisunla™), had been beforehand approved by the FDA, as demonstrated the elimination of Aβ plaques in the brains of individuals with Alzheimer’s disease (via surrogate markers) and improved some cognitive measures [10,11]. Nonetheless, some controversy remains regarding the overall efficacy and safety of these treatments [12–15].

A key aspect in addressing these diseases is the identification of reliable biomarkers, as they improve diagnostic accuracy, enable early detection, and are essential for developing and monitoring disease-modifying treatments. Ongoing research and standardization efforts aim to integrate these biomarkers into routine clinical practice, ultimately enhancing disease management and offering patients better therapeutic opportunities [16–19].

In the recent years, the development of biomarkers for neurodegeneration has rapidly advanced, including neuroimaging biomarkers and fluid-based biomarkers. Of particular importance are the cerebrospinal fluid (CSF) biomarkers. The underlying principle is simple: since CSF fills both the brain ventricles and the subarachnoid space, and exchanges substances with the brain and spinal cord, it can potentially reflect the pathological processes occurring in these structures. Compared to blood, CSF is less prone to peripheral contamination, which theoretically should facilitate the identification and interpretation of potential biomarkers. Additionally, the concentration of key proteins of interest is higher in CSF than in blood [20].

Different biomarkers have been encountered for AD, which permit not only determine the disease at the symptomatic phase but also in the prodromal phase, particularly those regarding the amyloidogenic process [18,21]. In this context, and with a focus on CSF, the most studied biomarkers include the p-tau/Aβ_42_ and Aβ_42_/Aβ_40_ ratios, which have demonstrated high sensitivity, specificity, and overall diagnostic accuracy for detecting intermediate-high AD neuropathologic changes [22]. Furthermore, CSF p-tau231 has been found to increase early in the progression of AD pathology and has been proposed as a leading candidate biomarker for identifying incipient Aβ pathology in the context of therapeutic trials [16,23,24].

Although research on CSF biomarkers for AD is well advanced, the availability of reliable biomarkers for other neurodegenerative disorders remains limited. This is particularly true for FTLD, where detecting the water-insoluble TDP-43 and FUS proteins in CSF presents a major challenge [7]. In this context, advances in precise analytical techniques, such as Single Molecule Array (Simoa) technology and Seed Amplification Assays, which enable the detection of extremely low levels of specific components in the CSF, can be particularly useful [20,25,26,27]. These methods will also enable the detailed characterization of these components, including aspects such as isotypes or post-translational modifications, which are particularly valuable in the context of protein aggregation [27].

At this point, it is worth highlighting that CSF studies often aim to detect proteins with very low solubility in aqueous media, particularly those that tend to aggregate in various proteinopathies. These include, among others, the above mentioned Aβ, tau, α-synuclein, FUS and TDP-43. Moreover, some of these proteins can propagate in a prion-like manner, and thus misfolded proteins can act as a seed to disrupt the normal proteins and produce their aggregation and precipitation. Accordingly, this seeding process can probably limit the presence of the free protein fraction in the CSF.

Given this context, a critical fact deserves special attention and is highly relevant to our point of view. In standard CSF studies, samples are routinely centrifuged, and the sediments discarded to eliminate cellular debris and high-density particles [28,29,30]. Thus, the question proposed is simple: may the discarded sediments contain components that could be useful as biomarkers, such as the insoluble amyloid fibrils or other aggregate-prone proteins, that tend to aggregate and precipitate? Exploring this possibility could open new avenues for biomarker discovery in neurodegenerative diseases.

## Results

To address whether the CSF sediments may contain components useful as biomarkers, a set of experiments detailed below were conducted. All experiments were performed with brain and CSF samples from two specific cases: one donor with AD and another with FTLD-Tau, specifically with progressive supranuclear palsy (PSP). The diagnosis and disease progression of both donors were confirmed post-mortem through neuropathological examination. Demographic and neuropathological data are provided in Supplementary Table 1. The experiments included, among others, the study of the ultrastructure of the CSF pellets with both transmission and scanning electron microscopy techniques (TEM and SEM respectively). Additionally, SEM combined with energy-dispersive X-ray spectroscopy (EDX) was used to obtain complementary information about the composition of the different structures observed in the pellets. Moreover, semithin sections measuring 500 nm in thickness were obtained and processed, following an adapted electron microscopy procedure, to apply chemical and immunofluorescence stains in order to obtain more precise information about their composition. All these procedures are detailed in the Methods section.

### Representative brain features of the AD and PSP donors

Based on the neuropathological diagnosis, we first performed indirect immunostaining for Aβ and tau on specific hippocampal sections from both donors. This allowed to visualize the representative brain features of each case in terms of tau and Aβ deposits, and to validate the antibodies that would later be used for staining the CSF pellets from both donors. For the Aβ staining, we used the 6E10 antibody, directed against several isoforms of Aβ, and the 12F4 antibody, specifically targeting the Aβ_42_ isoform. For tau staining, we used the Tau5 antibody, which recognizes various isoforms of tau. Preadsorption procedures were conducted to confirm the specificity of these stains, and negative controls were performed by incubating the samples with the buffer solution instead of the primary antibody.

All these stains were combined with that of the p62 protein, which is a well-established marker of wasteosomes [31,32]. It is worth noting that, as will be discussed later, wasteosomes are bodies that accumulate in the human brain and can be thereafter released into the CSF [32]. Therefore, it is expected to detect them both in the hippocampal sections and in the CSF pellets that will subsequently be analysed.

As expected and illustrated in Figure 1, the hippocampus of the AD patient contains Aβ plaques that become visible with both the 6E10 and 12F4 staining, and also contains the characteristic neurofibrillary tangles (NFTs), which become visible with the Tau5 staining. In the PSP sections, the 6E10 and 12F4 stains do not appear because of the absence of Aβ plaques in its hippocampus, while Tau5 permits to observe both the characteristic globose NFTs and tau-positive tufted astrocytes. In all cases, the preadsorption of the primary antibodies with the respective targeted protein abolished the staining. Moreover, and as expected (although not shown in the figure) wasteosomes are present in the periventricular and subpial regions of the hippocampus in both donors, and they become stained by the p62 staining. The p62 antibody also reacts with NFTs present in the AD hippocampus, and thus some NFTs become double stained with both the p62 and the Tau5 antibodies.

**Figure 1.**
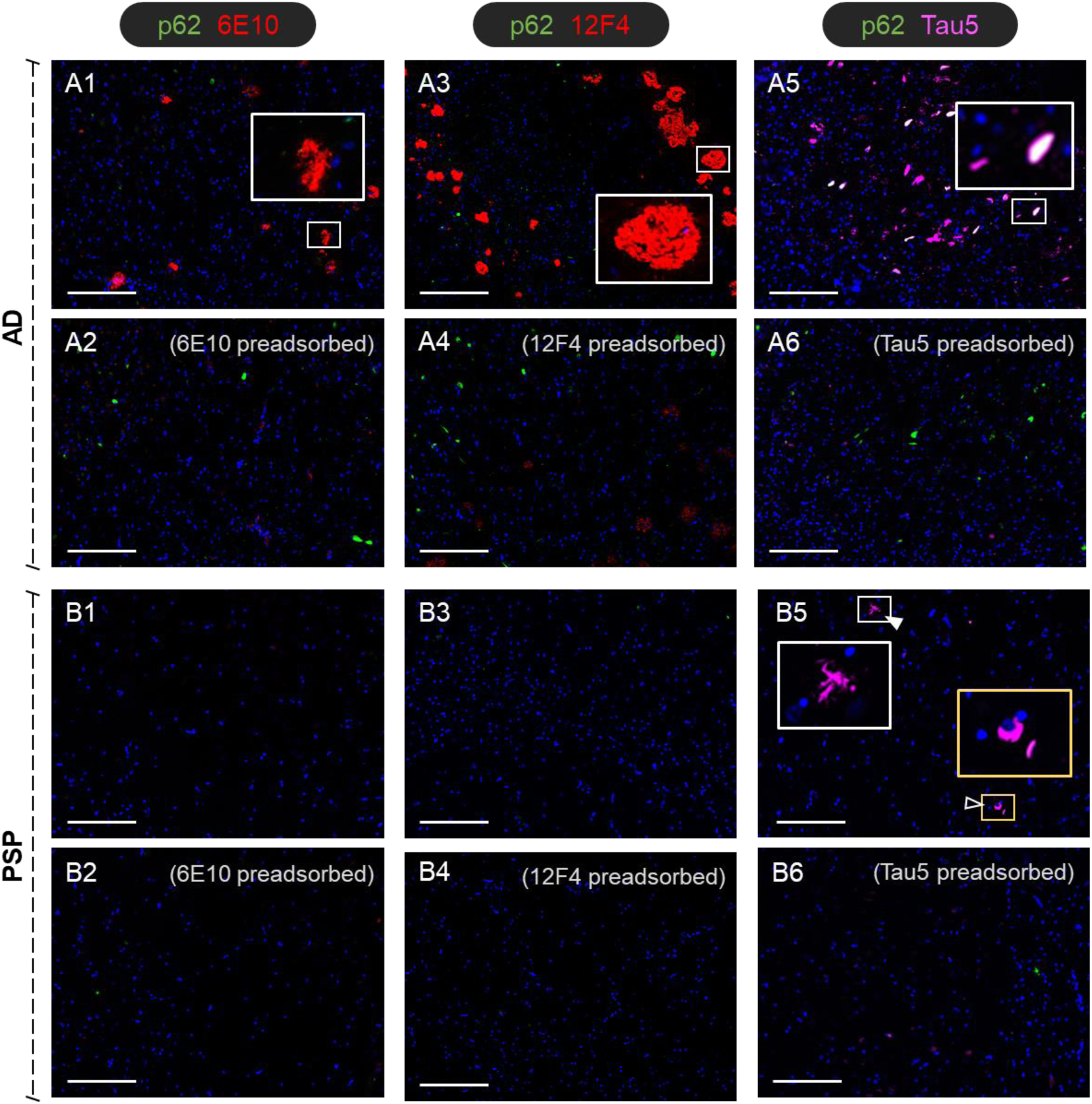
Representative images of the CA1 region of hippocampal sections from the AD donor (**A1-A6**) and the PSP donor (**B1-B6**) double immunostained with anti-p62 and 6E10, anti-p62 and 12F4 or anti-p62 and Tau5. In some images, some insets are magnified in order to facilitate the visualisation of specific structures. The AD patient showed the characteristic Aβ plaques stained with 6E10 and 12F4 (**A1**, **A3**), whereas the PSP patient did not display these plaques (**B1**, **B3**). Preadsorption of both antibodies with Aβ42 peptide (**A2**, **A4**, **B2**, **B4**) abolished the staining observed in the AD case. Regarding tau, the AD case exhibited abundant and widespread NFTs, some of which can be observed in white colour because they become also stained with the p62 (**A5**). The staining with Tau5 in the PSP case showed the characteristic tufted tau-positive astrocytes and globose-shaped NFTs (**B5**, arrowhead and empty arrowhead respectively). Tau staining disappeared in both AD and PSP cases after pre-adsorption of Tau5 with tau protein (**A6**, **B6**). All sections contain Hoechst staining (blue). Scale bars: 200 µm.

### A first inspection of the CSF pellets

For a preliminary inspection of the CSF pellets, we stained some semithin sections from each pellet with the periodic acid-Schiff (PAS) technique. To obtain the semithin sections, the intraventricular CSF was centrifuged, the supernatant eliminated, and the pellet embedded in resin to obtain the 500 nm-thick sections thereafter. It should be noted that, in the present study, PAS staining was performed on semithin sections; therefore, the staining intensity is reduced compared to that observed in standard histological sections.

As illustrated in Figure 2, the staining of both the AD and PSP samples revealed scattered, lightly stained components alongside structures with more intense staining. Among these structures, wasteosomes were identified, some of them within the pellet deposits and others at the edges of the semithin sections, likely due to redistribution during the centrifugation step of sample preparation. Additionally, in the pellets from both patients, as well as in additional cases not included in this study, PAS staining revealed faintly stained structures consistent with psammoma bodies.

**Figure 2.**
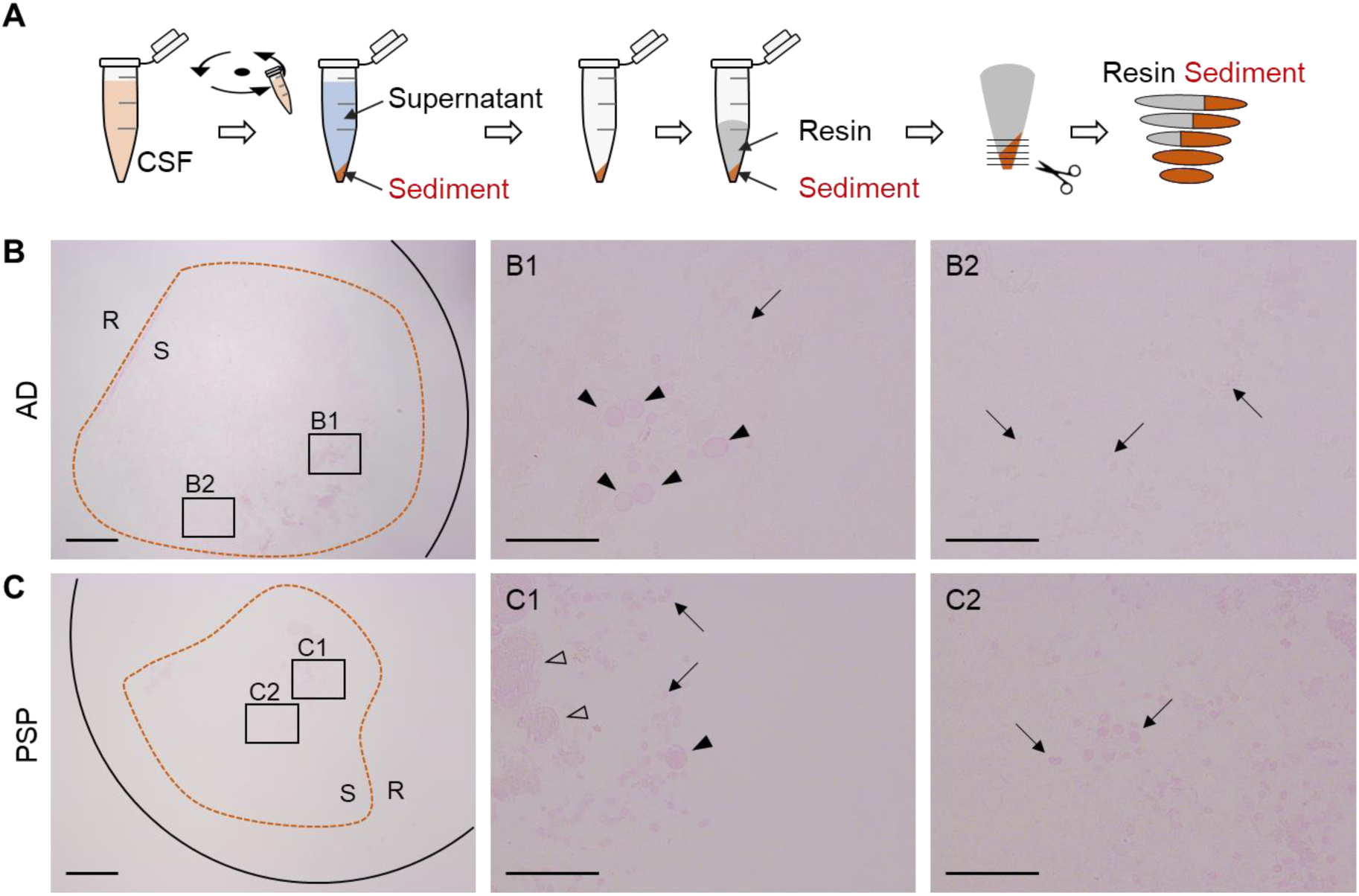
Representative images of semithin sections obtained from CSF pellets and stained with PAS. **A)** To obtain the semithin sections, CSF samples are centrifuged, the supernatant is then eliminated, and the pellet is resin embedded in order to obtain the sections (500 nm-thick). **Β-C)** Representative images of the PAS-stained semithin sections obtained from the AD and the PSP donors. These sections include a sediment area (S), as well as a region containing only resin (R). As can be observed in the insets, the sediments can contain wasteosomes (arrowheads), and psammoma bodies (empty arrowheads), alongside other unidentified pellet deposits (arrows). Scale bars on B and C: 400 µm. Scale bars on insets: 100 µm.

As commented before, the presence of wasteosomes in the CSF and their brain origin had already been described [32], and serves as evidence of the presence of brain-derived components in these pellets. On the other hand, the presence of psammoma bodies in CSF pellets, which will be confirmed with the following tests, had not been previously described.

### Ultrastructural characterization of CSF pellets using TEM

Based on the observations made using the PAS staining technique, certain regions of interest (ROIs) have been identified in the different semithin sections, and ultrathin sections of these ROIs have been obtained and processed for visualization under TEM.

As illustrated in Figure 3 for the PSP case, the TEM analysis of the ultrathin sections of the CSF pellets from both donors revealed not only a variety of amorphous structures, but also membranous debris, fibrillar structures and vesicles, as well as wasteosomes and psammoma bodies. As indicated before, the presence of wasteosomes in the CSF and their brain origin had already been described, but not that of the psammoma bodies. To investigate further, we analysed haematoxylin and eosin-stained, paraffin-embedded sections from the temporal lobe and spinal cord of the studied donors, sections previously used in the donors’ neuropathological examinations, and observed the presence of both wasteosomes and psammoma bodies in both structures, located either within their parenchyma or in their bordering regions (Supplementary Figure 1). These observations support the brain origin of the psammoma bodies found in the CSF and strongly reinforce the presence of brain-derived components in the CSF pellets.

**Figure 3.**
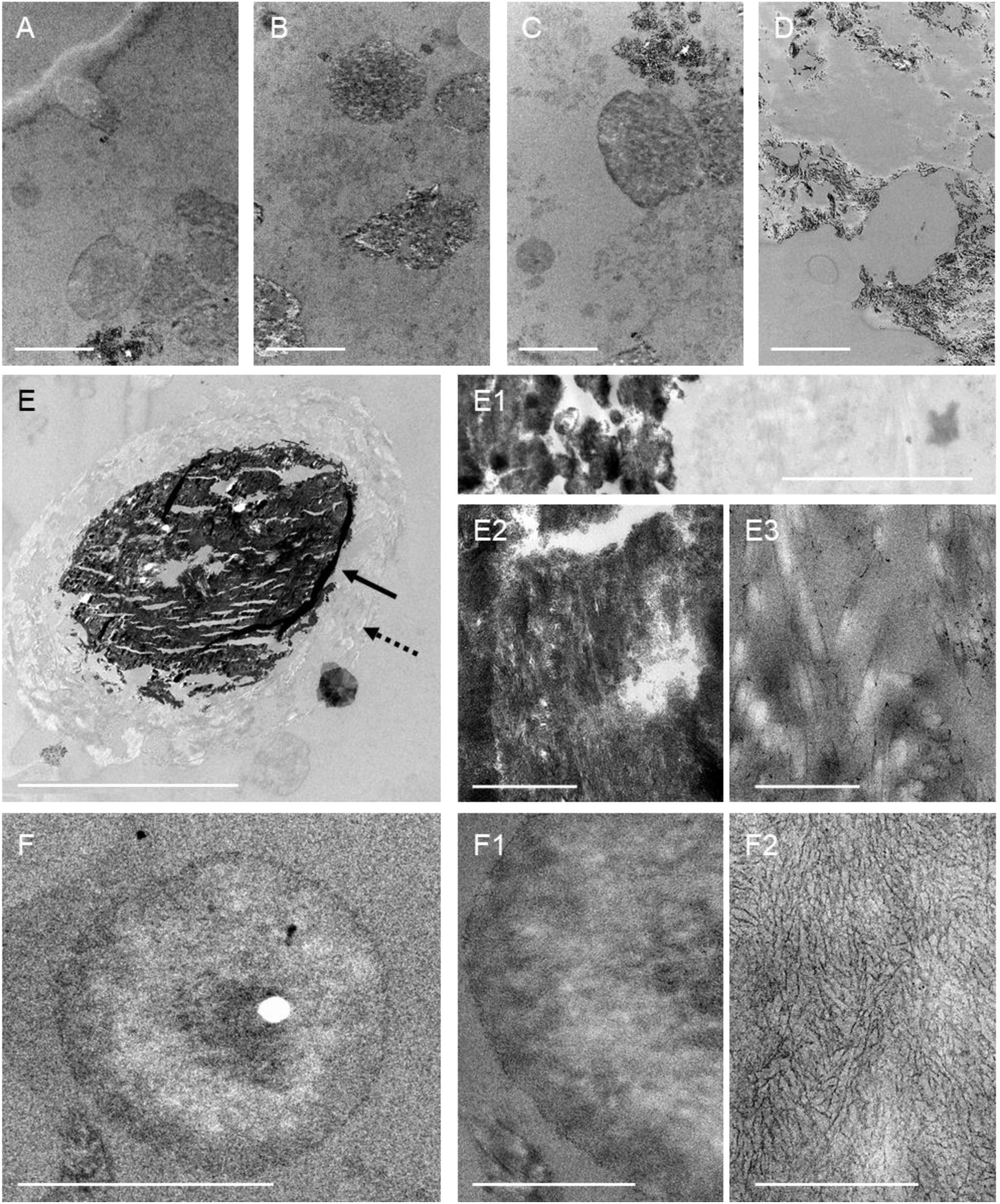
Representative TEM images of the CSF pellet obtained from the PSP case. **A-D)** TEM images highlighting amorphous structures and various remnants in different regions of the sample. Scale bars: 5 µm. **E)** Among the different structures, some of them are compatible with psammoma bodies. In this case, the psammoma contains a central electrodense region (arrow) and a peripheral non-electrodense region (dashed arrow). Scale bar: 20 µm. **E1)** Higher-magnification TEM image that includes the electrodense region of the psammoma body and its peripheral region. Scale bar: 2 µm. **E2-E3)** At higher magnification, fibrillar structures appear in both, the central and the peripheral regions of the psammoma. Scale bars: 500 nm. **F)** Representative TEM image showing a wasteosome. Scale bar: 20 µm. **F1-F2)** Higher-magnification TEM images showing the detailed ultrastructure of the wasteosome. As can be observed in F2, wasteosomes encompass densely packed, randomly oriented, short linear fibres. Scale bars: 20 µm (F1) or 500 nm (F2).

### Characterization of CSF pellets using SEM-EDX

Following TEM analysis, complementary observations were made using SEM imaging and SEM-EDX analysis. As expected, the SEM images permit to identify the presence of wasteosomes and psammoma bodies along with some unidentified remnants in the pellets from both the AD and the PSP patients. Figure 4 illustrates the results of these studies for the PSP case, and shows the presence of both types of structures (psammoma bodies and wasteosomes) in representative sections stained with PAS, the magnifications of some of these structures in consecutive sections observed on SEM, and some of the spectra obtained with the analysis performed by SEM-EDX.

**Figure 4.**
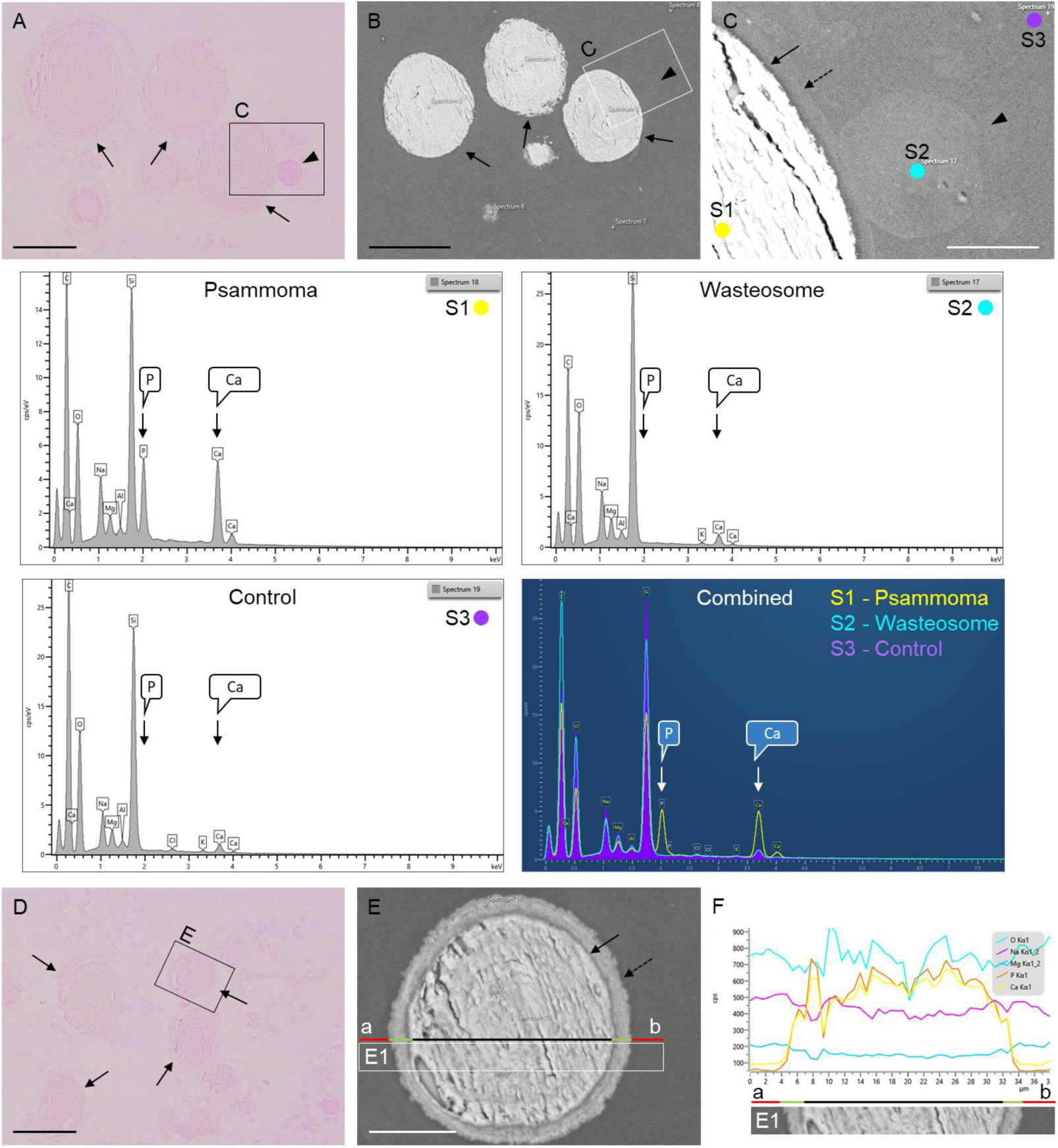
Representative images of SEM and SEM-EDX analyses of the CSF pellet obtained from the PSP case. **A**) PAS stained semithin section in which a wasteosome and some psammoma bodies can be observed (arrowhead and arrows respectively). **B**) SEM image of the equivalent region, obtained in a consecutive section. **C**) SEM image corresponding to the C area marked in A and B. The dashed arrow indicates the peripheral region of a psammoma, the black arrow its core, and the arrowhead indicates a wasteosome. The SEM-EDX spectra obtained in the **S1**, **S2**, and **S3** points (corresponding respectively to the psammoma, wasteosome, and control regions) are shown below. Note the high levels of P and Ca on the psammoma body, clearly visible on the combined graph. **D**) PAS stained semithin section showing various psammoma bodies (black arrows). The psammoma body included in the area signalled as E was further analysed by SEM-EDX. **E**) SEM image of this psammoma body, obtained in a consecutive slide. The dashed arrow indicates its peripheral region and the black arrow indicates its core. **F**) Profile of the SEM-EDX analysis along the a-b line indicated in E. Note that in the core of the psammoma (line in black), the levels of P and Ca are increased when compared with that of the control region (line in red). The levels of such components show intermediate values in the peripheral region of the psammoma (line in green). E1 corresponds to the inset signalled in E. Scale bars: 50 µm (in black) or 10 µm (in white).

In the SEM-EDX analysis, we observed that the spectrum of the wasteosomes does not differ from that of the surrounding region (control region). It is important to note that the main components of wasteosomes are carbohydrates, and therefore, a high presence of carbon (C), hydrogen (H), and oxygen (O) would be expected. Nonetheless, the detection of carbon is masked by the carbon coating required for analysing non-conductive samples. In the case of O, it can be seen that the control region shows a high presence of silicon (Si) and O due to the glass slide, and this O masks the signal from the O present in the wasteosomes. On the other hand, hydrogen, due to its low molecular weight, cannot be reliably detected by SEM-EDX. In contrast, SEM-EDX analysis did reveal significant differences between psammoma bodies and the control regions. As shown in Figure 4, there is a high presence of calcium (Ca) and phosphorus (P) in the core of the psammoma bodies, supporting their identification as such.

### Immunofluorescence on CSF pellets

After confirming the presence of both wasteosomes and psammoma bodies in the CSF, we studied the presence of other components using immunofluorescence on semithin sections.

To begin with, we performed particular immunostainings based on previous studies. Specifically, we previously observed that after resuspending the pellet and spreading it onto a slide, wasteosomes present in CSF can be immunostained with antibodies α-ubiquitin (Ubi), α-p62 (p62), α-glycogen synthase (GS) and natural IgMs (IgMs) [32]. Therefore, as a first step, we tested whether these immunostainings could also be observed in wasteosomes sectioned in the semithin slices used for our analyses. As illustrated in Figure 5, results indicate that wasteosomes on semithin sections from both donors indeed exhibit a well-defined positive labelling with Ubi, p62, and IgMs. In contrast, GS staining was very weak. As expected, the staining with Ubi, p62, and IgMs appears in the peripheral regions of the wasteosomes, showing a well-defined circular or ring-like structures with uniform staining. Moreover, some wasteosomes also present a central core that becomes stained with Ubi, but not with the other antibodies. The staining of GS, although faint, is also located in the peripheral regions of the wasteosomes. Altogether, these observations validate the use of immunofluorescence techniques on these semithin sections. Moreover, it is important to highlight that, in addition to the labelling observed in the wasteosomes, there is also some labelling in other regions of the pellets (also illustrated in Figure 5). Thus, staining with IgMs reveals numerous circular deposits scattered across the field, often accompanied by smaller, punctate, and less organized structures. Ubi staining exhibits a similar pattern to that observed with natural IgMs. Likewise, p62 staining displays diffuse granular patterns. Notably, no GS staining is observed in the pellet deposits. The presence of such components that become stained with Ubi, p62, and IgMs is coherent with the presence of remnants in the pellet, as all these stains are related to the signalling and removal of waste products [31–33].

**Figure 5.**
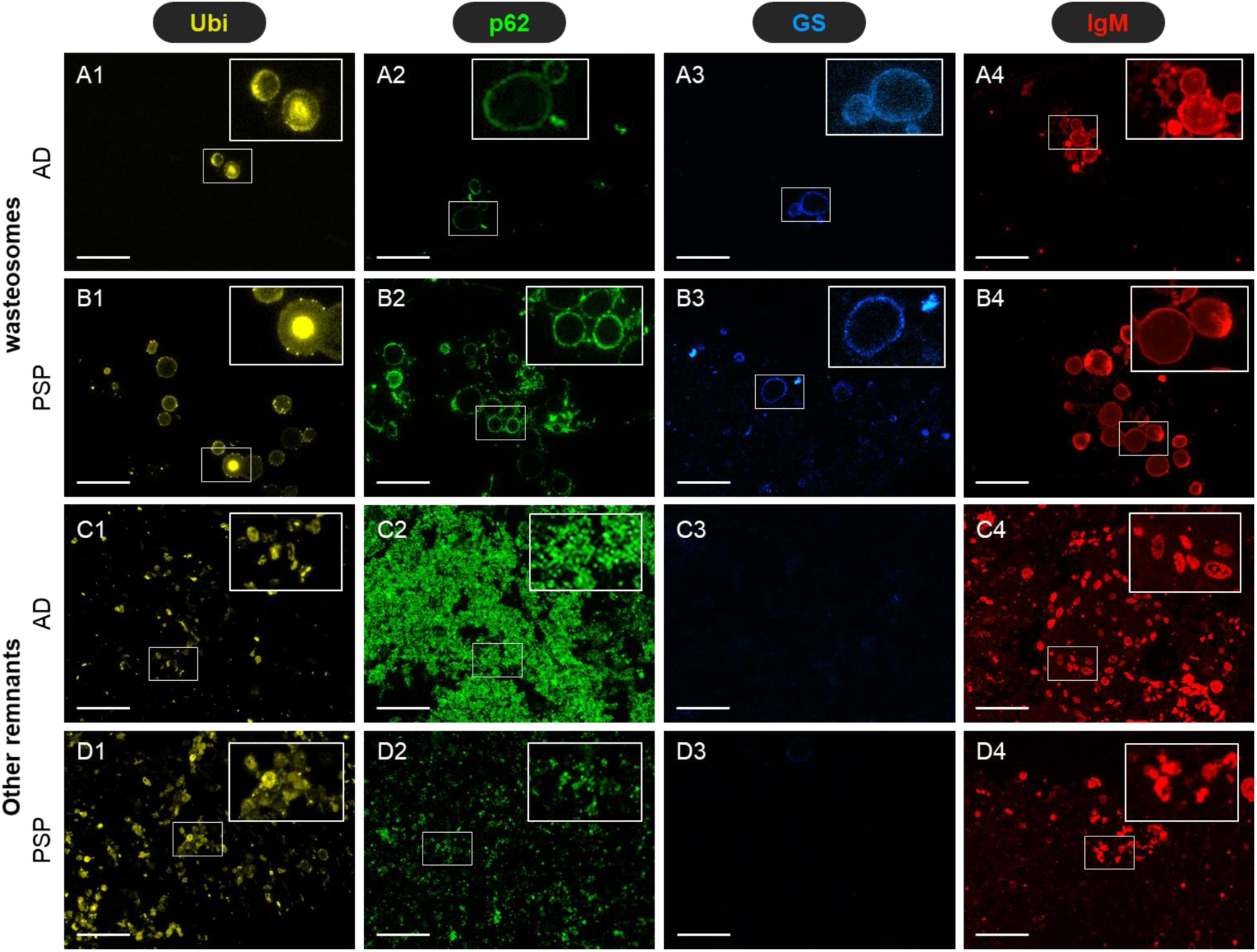
Representative images of semithin sections of CSF pellets obtained from the AD and PSP donors immunostained with the indicated antibodies. **A1**-**A4)** Images from the AD donor showing the stainings of the wasteosomes with Ubi, p62, GS, and IgMs respectively. One region of each image (inset) is magnified and colour adjusted to facilitate its visualisation. **Β1**-**B4)** In this case, the images correspond to the PSP donor. **C1**-**C4)** and **D1**-**D4)** illustrate the presence of stain in other regions of the pellet, in the AD and PSP sections respectively. Note the stainings of the wasteosomes in the sections from both donors, and the presence of some remnants that become stained by Ubi, p62, and IgM. Scale bar: 50 µm.

### Presence of disease-related proteins in CSF pellets

Given the remnants observed with the different techniques both in the ultrathin and the semithin sections of the CSF pellets, and their potential to contain information overlooked by the standard techniques commonly used, we further studied the presence of some disease-related proteins among these remnants. As it is well known, both Aβ and tau proteins are affected in the case of AD, while only the second one is affected in PSP. Accordingly, we performed 6E10, 12F4, and Tau5 immunostainings on semithin sections, adding both the preadsorption experiments to verify the specificity of the stainings and the respective controls by incubating the samples with BB instead of the primary antibodies. As mentioned earlier, the 6E10 and 12F4 antibodies are directed against Aβ and the isoform Aβ_42_ respectively, while Tau5 is directed against the tau protein.

Notably, immunostaining with 6E10 and 12F4 revealed that the semithin sections from the AD patient, but not those from the PSP patient, contain irregular and amorphous structures labelled by both antibodies. As these stainings disappear when the primary antibodies are preadsorbed with Aβ_42_, these findings denote the presence of Aβ_42_ in the pellets of the AD donor. On the other hand, the immunostaining with Tau5 revealed the presence of tau in the pellets from both the AD and the PSP donors, with particularly strong labelling in the AD donor. During the preadsorption studies with tau protein, the disappearance of the tau labelling was evident in both the AD and PSP, although some background or nonspecific labelling was noted on PSP. The positive stainings and the preadsorption results are summarized in Figure 6. Altogether, these results indicate that CSF pellets from AD contain both Aβ and tau proteins, whereas those from PSP contain only tau protein, remarkably reflecting the characteristic protein profiles of each disease.

**Figure 6.**
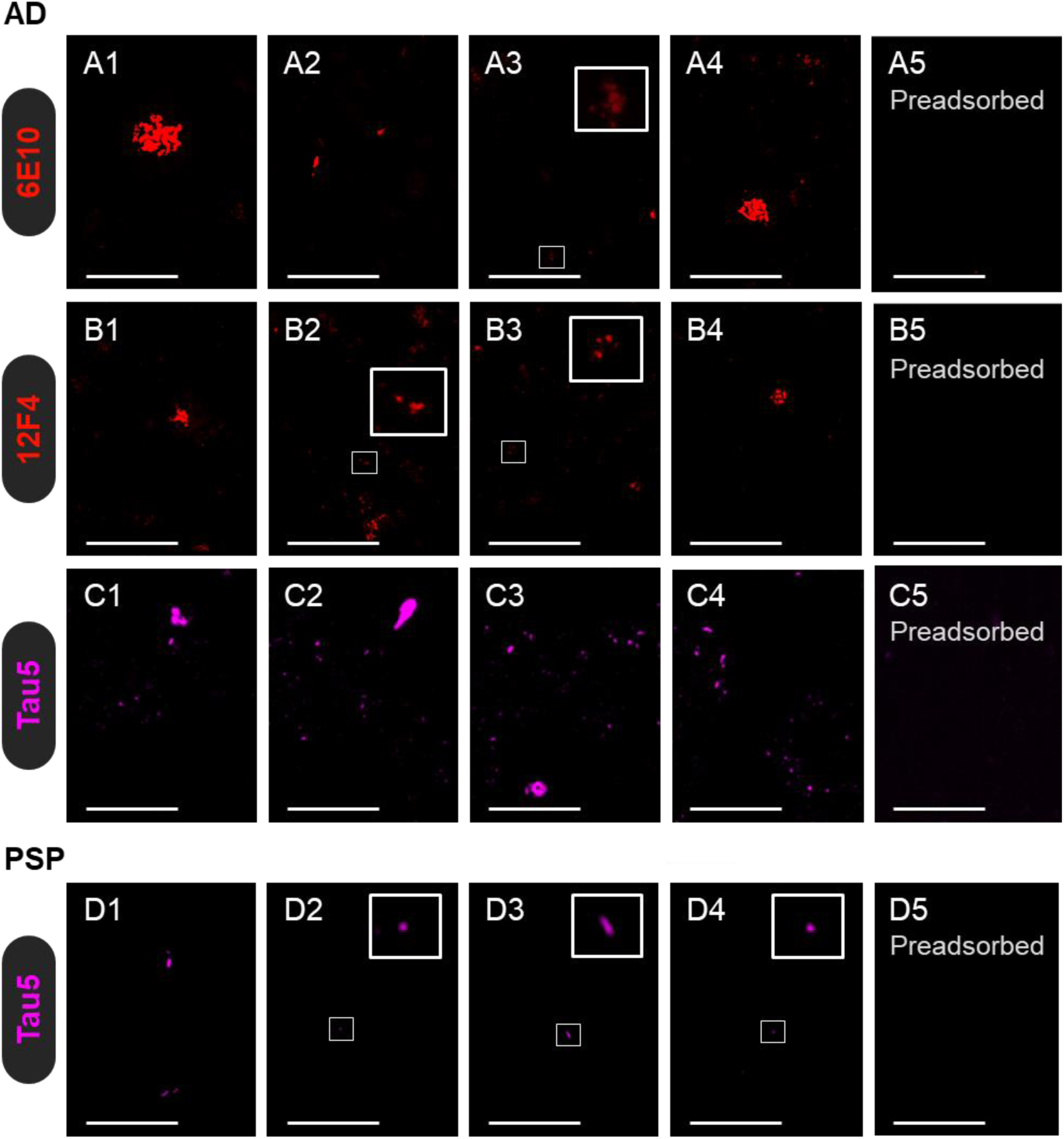
Representative images of semithin sections of CSF pellets obtained from the AD and PSP donors immunostained with 6E10, 12F4, and Tau5. Note that in some images some insets are magnified and colour adjusted to facilitate the visualisation of specific components that otherwise remain unappreciated. **A1-A4)** Sections from the AD donor immunostained with 6E10 show varied positive structures. **A5)** When preadsorbing the 6E10 with Aβ42 the staining does not appear. **B1-B6)** The same procedure but with the 12F4 antibody produces similar results: some components become immunostained, and the preadsorption of the 12F4 antibody produces the absence of staining. **C1-C6)** When using Tau5, some tau positive structures can be observed, and the preadsorption with the tau protein prevents such staining. **D1-D6)** For the PSP donor, the staining with Tau5 also produces the staining of some structures, which mainly disappear when the antibody is preadsorbed with the tau protein. The 6E10 and 12F4 antibodies did not produce any staining on the PSP sections (images not shown). Altogether, these results indicate that CSF pellets from AD contain both Aβ and tau proteins, whereas those from PSP contain only tau protein, remarkably reflecting the characteristic protein profiles of each disease. Scale bar: 50 µm.

## Discussion

The evidence gathered in the present study highlight that potential biomarkers may accumulate or be hidden within CSF sediments. The findings indicate that the CSF pellet from the AD case contains Aβ, particularly Aβ_42_, which has very low solubility and tends to aggregate. Additionally, tau protein can also be contained in CSF pellets, as evidenced by both the significant tau labelling observed in the semithin sections from the AD donor and the faint staining observed in the PSP donor. As the results are based on samples from only two patients, we cannot extrapolate that the CSF sediments of all patients will contain these proteins. However, we can affirm that CSF sediments may contain proteins of interest, which is already relevant in itself and constitutes a first step in this area of research. Furthermore, the fact that the proteins that have been detected in the semi-thin sections of the CSF sediment correspond to the altered proteins of each disease reinforces the interest of its study.

At this point, it should be noted that CSF samples used in the study are intraventricular fluid samples obtained postmortem. Accordingly, the study needs to be expanded towards the presence of these proteins in the pellets of CSF samples obtained by lumbar puncture in living people. It is expected that these samples will also present these proteins in the solid fraction or sediment, because it is well known that their liquid fraction contains them. Moreover, it should be noted that material exhibiting some characteristics of Aβ had previously been noted in CSF sediments, based on observations made in electron microscopy and staining of the sediment with Thioflavin S [34]. Furthermore, in studies that we are carrying out, we observe that CSF samples obtained by lumbar puncture contain wasteosomes, and therefore we know that their solid fraction or sediment contains components of cerebral origin.

Regarding Aβ in the CSF, it is widely accepted that AD patients exhibit a reduction in Aβ_42_ on the supernatant fraction of the CSF, with the prevailing hypothesis being that Aβ aggregation and retention into plaques in the brain parenchyma reduces its diffusion into the CSF [35]. Our results suggest an alternative or complementary explanation: significant amounts of Aβ may still be present in CSF but are sequestered in the sediment following centrifugation, rather than remaining in the supernatant. The transformation of soluble Aβ monomers into insoluble monomers and fibrillar aggregates that contain Aβ in a β-pleated sheet conformation is described as “seeded polymerization”. From our point of view, it cannot be ruled out that this process not only takes place in the brain parenchyma, inducing the formation of amyloid plaques, but also takes place in the CSF itself. Actually, the principle of seeded polymerization combined with fluorescence correlation spectroscopy permits the detection of amyloid β-peptide aggregates in CSF samples from AD patients [36], and liquid-based atomic force microscopy has revealed that the morphology of protein aggregates in the CSF differs distinctly between AD and non-AD patients [37]. Remarkably, in the present study, despite using a relatively low-sensitivity detection method, we were still able to identify both Aβ and tau in CSF sediments, suggesting that their presence may be substantial.

Additionally, because these aggregates in the CSF must be relatively large, they are likely to be cleared through the meningeal lymphatics and be phagocytosed by meningeal macrophages or transported to the cervical lymph nodes and beyond [38–41]. The efficient clearance of such protein aggregates and other waste products from the brain relies on the coordinated action of the glymphatic system, which facilitates fluid movement across the brain parenchyma, and the meningeal lymphatic system [40, 42–45]. Notably, a recent study indicates that fluid biomarkers are enriched in human cervical lymph nodes, with tau levels in lymph fluid being 266 times higher than in blood [46]. This finding aligns with our observations, suggesting that tau and other biomarkers may be present in CSF at higher concentrations than previously detected, potentially due to losses during the centrifugation process.

In addition to disease-related proteins, our results indicate that CSF pellets may also contain wasteosomes and psammoma bodies. As previously mentioned, the presence of wasteosomes in CSF pellets from patients with AD has already been reported [32]. Wasteosomes, originally known as corpora amylacea [47], are brain structures whose quantity increases with age and is further elevated in the context of various neurodegenerative diseases [47–56]. Proposed to be a hallmark of chronic glymphatic insufficiency [57], their function is unclear, although it has recently been proposed that they can act as containers that remove waste products from the brain [32]. In fact, they can be released from the brain into the CSF [32,58], and from there, they can be transported to the cervical lymph nodes via the meningeal lymphatic system [32]. Additionally, they have been shown to undergo phagocytosis by macrophages in vitro [32, 59, 60].

Consistent with their proposed role as waste containers, several studies have shown that wasteosomes from the brains of individuals with certain diseases can contain proteins associated with those diseases. For example, in the brain of patients with AD, wasteosomes may contain tau protein, although not Aβ [52,53], while in cases of FTLD, they may harbour traces of tau, FUS, or TDP-43, depending on the FTLD subtype [54]. Based on these findings, it would be reasonable to expect that some of the wasteosomes observed in the CSF pellets might contain tau protein in the AD cases, and possibly in the PSP cases as well. However, in the sections we analysed, we were unable to detect tau presence within the wasteosomes. It should be noted that we were working with semithin sections of CSF pellets, and it cannot be ruled out that the amount of protein present in the sliced wasteosomes was below the detection threshold for immunofluorescence. Moreover, the presence of tau in the wasteosomes found in the brains of disease-affected donors was not a general feature of all wasteosomes, but rather only some of them contained the protein.

In reference to psammoma bodies, their presence in the CSF pellets has not previously been reported. Psammoma bodies are lamellar, calcified structures commonly found in both neoplastic and non-neoplastic conditions. They typically measure between 50 and 150 μm in diameter and are primarily composed of calcium in the form of phosphate and hydroxyapatite, although trace elements such as iron have also been detected [61,62]. These structures have been particularly associated with meningiomas, age-related changes, and epithelial atrophy [63–66]. In the 19th century, Virchow noted that psammoma formations are “an everyday occurrence in the region of the medulla oblongata where they appear in the nipple of choroid plexus on the roof of the fourth ventricle” [63]. Given that the choroid plexus serves as the primary site of CSF production and secretion, the detection of psammoma bodies in CSF pellets is thus not unexpected. Moreover, the presence of psammoma bodies is not only restricted to this choroid plexus related-region. As indicated, we also observed them in the temporal lobe and spinal cord, located either within their parenchyma or in their bordering regions, and we do not rule out their presence in other unchecked brain regions. This evidence also reinforces the brain origin of the psammoma bodies observed in the CSF pellets.

In conclusion, the evidence gathered in this study indicates that CSF pellets contain several identifiable brain-derived components, including wasteosomes, psammoma bodies, and deposits of disease-related protein aggregates that closely reflect the patient’s underlying proteinopathy. Consequently, future biomarker research should consider analysing these sediments or adapting existing methodologies to study uncentrifuged CSF directly. Investigating CSF pellets or uncentrifuged CSF may offer valuable insights into the pathophysiology of neurodegenerative diseases, ultimately improving early diagnosis and disease monitoring.

## Methods

### Human brain samples

Post-mortem cryopreserved hippocampal sections (6 µm-thick) were obtained from one neuropathologically confirmed case of AD with Down Syndrome and one from FTLD-PSP. The neuropathological assessment followed standardized protocols established by the Neurological Tissue Bank (Biobank-Hospital Clínic-IDIBAPS, Barcelona) as well as international consensus criteria [67]. Briefly, half of each donor’s brain was fixed in a 4 % formaldehyde solution for three weeks, after which paraffin-embedded sections (5 µm-thick) were prepared from at least 25 brain regions to support the diagnosis. The other half of the brain was freshly dissected. A fragment of the hippocampus was collected and immersed in 4 % paraformaldehyde (4 °C) for 24 h, followed by a 48-h immersion in 30 % sucrose in phosphate-buffered saline (PBS) (4 °C). Once dried, the hippocampal fragment was frozen at −20 °C and then stored at −80 °C. Subsequent cryostat sectioning at 6 µm enabled use in immunofluorescence and histochemical analyses. All procedures, including the preparation and storage of hippocampal samples, were conducted by the Neurological Tissue Bank (Biobank-Hospital Clínic-IDIBAPS, Barcelona).

Brain tissue and CSF samples (see below) were collected with written informed consent from patients or their legal representatives, permitting the use of brain tissue and medical records for research, as authorized by the Ethics Committee of the Neurological Tissue Bank of IDIBAPS Biobank, following the Declaration of Helsinki. Experiments involving human tissue complied with relevant ethical guidelines and regulations and received approval from the Bioethical Committee of the University of Barcelona (IRB00003099).

### Human CSF samples

Post-mortem intraventricular CSF samples were collected from the same donors from whom hippocampal sections were obtained. Samples were initially centrifuged at 1000 *g* for 5 min, and the supernatants were discarded. Pellets were then resuspended in 0.1 M Phosphate Buffer (PB) and centrifuged again at 700 *g* for 10 min, with supernatants discarded. This washing step was repeated twice. Next, pellets were fixed in 4 % paraformaldehyde in PB for 1 h with gentle agitation. Afterwards, pellets were centrifuged at 700 *g* for 10 min, and the supernatants were discarded. Pellets were stored at 4 °C and transported to the Electron Microscopy Unit (UME) at the Scientific and Technological Centres of the University of Barcelona (CCiTUB). They were then washed four times in deionized water for 10 min each, followed by dehydration in a graded ethanol series. Finally, pellets were embedded in EPON resin and polymerized in a heater. Semithin sections (500 nm-thick) were prepared from each sample using an ultramicrotome (EM UC7, Leica, Wetzlar, Germany) with a digital camera (IC90 E CMOS, Leica, Wetzlar, Germany).

### Immunofluorescence in brain samples

Hippocampal sections were air dried for 10 min at room temperature. For Aβ staining, samples were rehydrated in PBS for 5 min, incubated in formic acid at 70 % for 30 seconds, then briefly rinsed by immersion and washed in PBS for 5 min. For tau staining, sections were placed in a water bath set at 100 °C inside staining dishes containing citrate buffer (pH 6.0) for 7.5 min. The staining dishes were then removed from the water bath and left at room temperature for 20 min. After cooling down, the samples were washed with PBS. After the respective pre-treatments, hippocampal sections were blocked and permeabilized with 1 % bovine serum albumin (Sigma-Aldrich, Madrid, Spain) in PBS (blocking buffer, BB) containing 0.1 % Triton X-100 (Sigma-Aldrich) for 20 min. Samples were then washed with PBS and incubated for 21 h at 4 °C with the primary antibodies for double staining: a mouse monoclonal IgG_1_ against tau (Tau5), a mouse monoclonal IgG_1_ against Aβ_16_ (6E10) or a mouse monoclonal IgG_1_ against Aβ_42_ (12F4), and a mouse monoclonal IgG_2a_ against p62 (p62) (antibody details are compiled in Supplementary Table 2). Sections were then washed and incubated for 1 h at room temperature with the corresponding secondary antibodies: AF555 goat anti-mouse IgG_1_ (1:250; A-21121; Life Technologies; Eugene, OR, USA) and AF488 goat anti-mouse IgG_2a_ (1:250; A-21137; Life Technologies). Nuclei were then stained with the Hoechst stain (2 μg/mL; H-33258; Fluka; Madrid, Spain), and the samples were washed and coverslipped with Fluoromount (Electron Microscopy Sciences; Hatfield, PA, USA). Staining controls were performed by incubating with BB instead of the primary antibody before incubation with the secondary antibody.

Additionally, preadsorption studies were performed to verify the specificity of the staining. Specifically, anti-Aβ antibodies were incubated for 21 h at 4 °C with a 100-fold molar excess of synthetic antigen, then centrifuged at 16000 *g* for 30 min before being used for immunofluorescence. The same procedure was applied to the anti-tau antibody, with a 20-fold molar excess of synthetic antigen. Antigen data are summarized in Supplementary Table 2.

### PAS staining in CSF semithin sections

Semithin sections (500 nm-thick) were de-embedded by incubating in a 3:1 mixture of sodium methoxide and toluene/methanol for 5 min, followed by rinsing with toluene/methanol for 5 min, acetone for two 5-min washes, and a final water rinse. Subsequently, sections were stained using the PAS method, following the standard procedure previously described [33]. Briefly, the sections were fixed in Carnoy’s solution (60 % ethanol, 30 % chloroform, and 10 % glacial acetic acid) for 10 min. They were then pre-treated with 0.25 % periodic acid (19324-50, Electron Microscopy Sciences) in deionized water for 10 min, followed by a 3-min wash with deionized water. The samples were then incubated in Schiff’s reagent (26052–06, Electron Microscopy Sciences) for 10 min and washed for 5 min with deionized water. Nuclei were counterstained for 1 min with Mayer’s haematoxylin solution (3870, J. T. Baker). Afterward, the samples were washed, dehydrated with xylene, and mounted with Eukitt mounting medium (03989, Merck).

### Immunofluorescence in CSF semithin sections

Semithin sections (500 nm-thick) were de-embedded by incubating in a 3:1 mixture of sodium methoxide and toluene/methanol for 5 min, followed by rinsing with toluene/methanol for 5 min, acetone for two 5-min washes, and a final water rinse. For Aβ and tau stainings, semithin sections followed the same protocol as hippocampal sections, except for the water bath time in the tau pre-treatment, which was 20 min.

### Image acquisition

Immunohistochemistry and fluorescence images were captured using a fluorescence laser and optical microscope (BX41, Olympus, Hamburg, Germany) and stored in .tiff format. Exposure time was adjusted for each specific stain, with corresponding control images acquired using the same exposure settings. Image processing and analysis were conducted using the ImageJ software [68]. Any adjustments to contrast and brightness for improved visualization were applied uniformly to both the experimental and control images.

### TEM procedures

The semithin sections were first embedded in a Spur resin block, which was subsequently trimmed to a 200 µm^2^ area. Ultrathin sections (60 nm) were then obtained using an ultramicrotome (EM UC7, Leica, Wetzlar, Germany) with a digital camera (IC90 E CMOS, Leica, Wetzlar, Germany). The ultrathin sections were mounted on a 200-mesh copper grid and on a 200-mesh slot copper grid. Finally, they were stained with uranyl acetate for 4 min, followed by lead citrate for 2 min, and then washed.

The TEM analysis was conducted using a transmission electron microscope (JEM 1010 80 kV, JEOL, Tokyo, Japan) equipped with a side-mounted camera (Orius SC200D CCD, Gatan, Pleasanton, USA). Imaging was performed at an acceleration voltage of 80 kV.

### SEM-EDX procedures

The semithin sections were first coated with a carbon thin film to improve their electrical conductivity using a carbon evaporator (Q150Tplus, Quorum, Lewes, UK). Then, the sections were mounted to the holder using double-stick carbon tape and placed into the chamber. The SEM analysis was conducted using a Schottky-type field SEM (JSM-7100F, JEOL, Tokyo, Japan) equipped with an EDX system featuring a silicon drift detector Ultim Max (Oxford Instruments) and AztecLive as EDX software. The sample surface was examined at a magnification ranging from 120x to 3000x. Backscattered and secondary electron images were obtained at an acceleration voltage of 10 kV and a probe current of 10.

## Data availability

The authors declare that all data supporting the findings of this study are available within the paper and its supplementary information files.

## Acknowledgements

We are sincerely grateful to the Biobank-Hospital Clinic-Institut d’Investigacions Biomèdiques August Pi i Sunyer (IDIBAPS), integrated in the Spanish National Biobank Network, for providing the samples and data procurement. We are thankful to brain donors and families for generous brain donation for research. We thank Rosa Rivera and David Bellido from Scientific and Technological Centers of Universitat de Barcelona (CCiTUB) for their professional dedication, help and availability.

## Author contributions

Conception and Design: RA, MR, EP, JdV, CP and JV. Sample processing and execution: RA, MR, MS, CR, BV, EP, LMP, JdV, CP and, JV. Interpretation of data: RA, MR, EP, JdV, CP and JV. Writing and revising: RA, MR, BV, CP and JV. Critical review for important intellectual content: all authors. All authors have approved the final version of the manuscript.

## Funding

This work was supported by PID2020-115475GB-I00 and PID2023-148768OA-I00, both funded by MICIU/AEI/10.13039/501100011033, and the Generalitat of Catalonia (2021 SGR 00288 625). RA received the predoctoral Formación de Profesorado Universitario (FPU) 2022 fellowship from the MICIU. CR received the predoctoral Formación de Personal Investigador (FPI) 2022 fellowship from the MICIU. Hospital Clínic / FRCB-IDIBAPS receives support from the CERCA program of Generalitat de Catalunya. UBNeuro is supported by the Maria de Maeztu excellence centers program of the MICIU.

## Competing interests

The authors declare no competing interests.

## Notes

### Competing Interest Statement

The authors have declared no competing interest.

## References

1. Wilson, D.M. III. et al. Hallmarks of neurodegenerative diseases. Cell 186, 693–714 (2023).

2. Ross, C.A. & Poirier, M.A. Protein aggregation and neurodegenerative disease. Nat. Med. 10, S10–S17 (2004).

3. Soto, C. & Pritzkow, S. Protein misfolding, aggregation, and conformational strains in neurodegenerative diseases. Nat. Neurosci. 21, 1332–1340 (2018).

4. Knopman, D.S., et al. Alzheimer disease. Nat. Rev. Dis. Primers 7, 33 (2021).

5. Spillantini, M.G. et al. Alpha-synuclein in Lewy bodies. Nature 388, 839–840 (1997).

6. Mackenzie, I.R. et al. Nomenclature and nosology for neuropathologic subtypes of frontotemporal lobar degeneration: an update. Acta Neuropathol. 119, 1–4 (2010).

7. Grossman, M., et al. Frontotemporal lobar degeneration. Nat. Rev. Dis. Primers 9, 40 (2023).

8. GBD 2019 Dementia Forecasting Collaborators. Estimation of the global prevalence of dementia in 2019 and forecasted prevalence in 2050: an analysis for the Global Burden of Disease Study 2019. Lancet Public Health 7, e105–e125 (2022).

9. Brockmann, R., Nixon, J., Love, B.L. & Yunusa, I. Impacts of FDA approval and Medicare restriction on antiamyloid therapies for Alzheimer’s disease: patient outcomes, healthcare costs, and drug development. Lancet Reg. Health Am. 20, 100467 (2023).

10. Sadruddin, Z. et al. Anti-amyloid immunotherapies for Alzheimer’s disease: Administration, side effects, and overall framework. Geriatr. Nurs. 64, 103371 (2025).

11. Scheltens, P. et al. Alzheimer’s disease. Lancet 397, 1577–1590 (2021).

12. Terao, I. & Kodama, W. Comparative efficacy, tolerability and acceptability of donanemab, lecanemab, aducanumab and lithium on cognitive function in mild cognitive impairment and Alzheimer’s disease: A systematic review and network meta-analysis. Ageing Res. Rev. 94, 102203 (2024).

13. Daly, T., Kepp, K.P. & Imbimbo, B.P. Are lecanemab and donanemab disease-modifying therapies? Alzheimers Dement. 20, 6659–6661 (2024).

14. Espay, A.J., Kepp, K.P. & Herrup, K. Lecanemab and donanemab as therapies for Alzheimer’s disease: an illustrated perspective on the data. eNeuro 11, ENEURO.0319-23 (2024).

15. Høilund-Carlsen, P.F. et al. Donanemab, another anti-Alzheimer’s drug with risk and uncertain benefit. Ageing Res. Rev. 99, 102348 (2024).

16. Blennow, K., Hampel, H., Weiner, M. & Zetterberg, H. Cerebrospinal fluid and plasma biomarkers in Alzheimer disease. Nat. Rev. Neurol. 6, 131–144 (2010).

17. Molinuevo, J.L. et al. The clinical use of cerebrospinal fluid biomarker testing for Alzheimer’s disease diagnosis: a consensus paper from the Alzheimer’s Biomarkers Standardization Initiative. Alzheimers Dement. 10, 808–817 (2014).

18. Hansson O. Biomarkers for neurodegenerative diseases. Nat. Med. 27, 954–963 (2021).

19. Zetterberg, H. & Blennow, K. Moving fluid biomarkers for Alzheimer’s disease from research tools to routine clinical diagnostics. Mol. Neurodegener. 16, 10 (2021).

20. Alcolea, D., et al. Use of plasma biomarkers for AT(N) classification of neurodegenerative dementias. J. Neurol. Neurosurg. Psychiatry 92, 1206–1214 (2021).

21. Masters, C.L., et al. Alzheimer’s disease. Nat. Rev. Dis. Primers 1, 15056 (2015).

22. Mattsson-Carlgren, N. et al. Cerebrospinal fluid biomarkers in autopsy-confirmed alzheimer disease and frontotemporal lobar degeneration. Neurology 98, e1137– e1150 (2022).

23. Ashton, N. J. et al. Cerebrospinal fluid p-tau231 as an early indicator of emerging pathology in Alzheimer’s disease. EBioMedicine 76, 103836 (2022).

24. Cheslow, L., Snook, A.E. & Waldman, S.A. Biomarkers for managing neurodegenerative diseases. Biomolecules. 14, 398 (2024).

25. Scialò, C. et al. TDP-43 real-time quaking induced conversion reaction optimization and detection of seeding activity in CSF of amyotrophic lateral sclerosis and frontotemporal dementia patients. Brain Commun. 2, fcaa142 (2020).

26. Smith, R. et al. Misfolded alpha-synuclein in amyotrophic lateral sclerosis: Implications for diagnosis and treatment. Eur. J. Neurol. 31, e16206 (2024).

27. Teunissen, C.E. et al. Methods to discover and validate biofluid-based biomarkers in neurodegenerative dementias. Mol. Cell. Proteomics 22, 100629 (2023).

28. Teunissen, C.E. et al. A consensus protocol for the standardization of cerebrospinal fluid collection and biobanking. Neurology 73, 1914–1922 (2009).

29. Del Campo, M. et al. Recommendations to standardize preanalytical confounding factors in Alzheimer’s and Parkinson’s disease cerebrospinal fluid biomarkers: an update. Biomark Med. 6, 419–430 (2012).

30. Ashton, N.J. et al. Diagnostic accuracy of the plasma ALZpath pTau217 immunoassay to identify Alzheimer’s disease pathology. medRxiv [Preprint]. doi: 10.1101/2023.07.11.23292493 (2023).

31. Augé, E., Duran, J., Guinovart, J.J., Pelegrí, C. & Vilaplana, J. Exploring the elusive composition of corpora amylacea of human brain. Sci. Rep. 8, 13525 (2018).

32. Riba, M. et al. *Corpora amylacea* act as containers that remove waste products from the brain. Proc. Natl. Acad. Sci. USA. 116, 26038–26048 (2019).

33. Augé, E., Cabezón, I., Pelegrí, C. & Vilaplana, J. New perspectives on corpora amylacea in the human brain. Sci. Rep. 7, 41807 (2017).

34. Townsend, L.E. et al. Comparison of methods for analysis of CSF proteins in patients with Alzheimer’s disease. Neurochem. Pathol. 6, 213–229 (1987).

35. Fagan, A.M. et al. Inverse relation between in vivo amyloid imaging load and cerebrospinal fluid Abeta42 in humans. Ann. Neurol. 59, 512–519 (2006).

36. Pitschke, M., Prior, R., Haupt, M. & Riesner, D. Detection of single amyloid beta-protein aggregates in the cerebrospinal fluid of Alzheimer’s patients by fluorescence correlation spectroscopy. Nat. Med. 4, 832–834 (1998).

37. Nirmalraj, P.N., Schneider, T., Lüder, L. & Felbecker, A. Protein fibril length in cerebrospinal fluid is increased in Alzheimer’s disease. Commun. Biol. 6, 251 (2023).

38. Da Mesquita, S., Fu, Z. & Kipnis, J. The meningeal lymphatic system: a new player in neurophysiology. Neuron 100, 375–388 (2018).

39. Da Mesquita, S. et al. Functional aspects of meningeal lymphatics in ageing and Alzheimer’s disease. Nature 560, 185–191 (2018).

40. Jiang-Xie, L.F., Drieu, A. & Kipnis, J. Waste clearance shapes aging brain health. Neuron 113, 71–81 (2024).

41. Salvador, A.F.M., Abduljawad, N. & Kipnis, J. Meningeal lymphatics in central nervous system diseases. Annu. Rev. Neurosci. 47, 323–344 (2024).

42. Iliff, J.J. et al. A paravascular pathway facilitates CSF flow through the brain parenchyma and the clearance of interstitial solutes, including amyloid β. Sci. Transl. Med. 4, 147ra111 (2012).

43. Rasmussen, M.K., Mestre, H. & Nedergaard, M. The glymphatic pathway in neurological disorders. Lancet Neurol. 17, 1016–1024 (2018).

44. Holth, J.K. et al. The sleep-wake cycle regulates brain interstitial fluid tau in mice and CSF tau in humans. Science 363, 880–884 (2019).

45. Kipnis, J. The anatomy of brainwashing. Science 385, 368–370 (2024).

46. Al-Diwani, A. et al. Neurodegenerative fluid biomarkers are enriched in human cervical lymph nodes. Brain 148, 394–400 (2025).

47. Riba, M., Del Valle, J., Augé, E., Vilaplana, J. & Pelegrí, C. From corpora amylacea to wasteosomes: history and perspectives. Ageing Res. Rev. 72, 101484 (2021).

48. Averback, P. Parasynaptic corpora amylacea in the striatum. Arch. Pathol. Lab. Med. 105, 334–335 (1981).

49. Cissé, S., Perry, G., Lacoste-Royal, G., Cabana, T. & Gauvreau, D. Immunochemical identification of ubiquitin and heat-shock proteins in corpora amylacea from normal aged and Alzheimer’s disease brains. Acta Neuropathol. 85, 233–240 (1993).

50. Singhrao, S.K., Morgan, B.P., Neal, J.W. & Newman, G.R. A functional role for corpora amylacea based on evidence from complement studies. Neurodegeneration 4, 335–345 (1995).

51. Wilhelmus, M.M. et al. Novel role of transglutaminase 1 in corpora amylacea formation? Neurobiol. Aging 32, 845–856 (2011).

52. Wander, C.M. et al. The accumulation of tau-immunoreactive hippocampal granules and corpora amylacea implicates reactive glia in tau pathogenesis during aging. iScience 23, 101255 (2020).

53. Riba, M. et al. Uncovering tau in wasteosomes (*corpora amylacea*) of Alzheimer’s disease patients. Front. Aging Neurosci. 15, 1110425 (2023).

54. Alsina, R. et al. Increase in wasteosomes (corpora amylacea) in frontotemporal lobar degeneration with specific detection of tau, TDP-43 and FUS pathology. Acta Neuropathol. Commun. 12, 97 (2024).

55. Busard, H.L. et al. Polyglucosan bodies in brain tissue: a systematic study. Clin. Neuropathol. 13, 60–63 (1994).

56. Cavanagh, J.B. Corpora-amylacea and the family of polyglucosan diseases. Brain Res. Brain Res. Rev. 29, 265–295 (1999).

57. Riba, M., Del Valle, J., Molina-Porcel, L., Pelegrí, C. & Vilaplana, J. Wasteosomes (*corpora amylacea*) as a hallmark of chronic glymphatic insufficiency. Proc. Natl. Acad. Sci. USA. 119, e2211326119 (2022).

58. Sbarbati, A., Carner, M., Colletti, V. & Osculati, F. Extrusion of corpora amylacea from the marginal gila at the vestibular root entry zone. J. Neuropathol. Exp. Neurol. 55, 196–201 (1996).

59. Riba, M. et al. Corpora amylacea in the human brain exhibit neoepitopes of a carbohydrate nature. Front. Immunol. 12, 618193 (2021).

60. Riba, M. et al. Wasteosomes (corpora amylacea) of human brain can be phagocytosed and digested by macrophages. Cell Biosci. 12, 177 (2022).

61. Alcolado, J.C., Moore, I.E. & Weller, R.O. Calcification in the human choroid plexus, meningiomas and pineal gland. Neuropathol. Appl. Neurobiol. 12, 235–250 (1986).

62. Kubota, T., Sato, K., Yamamoto, S. & Hirano, A. Ultrastructural study of the formation of psammoma bodies in fibroblastic meningioma. J. Neurosurg. 60, 512–517 (1984).

63. Virchow, R. Die Krankhaften Geschwülste. 2, 106-113. Berlin: August Hirschwald (1863-1867).

64. Jovanović, I., Ugrenović, S., Antić, S., Stefanović, N. & Mihailović, D. Morphometric and some immunohistochemical characteristics of human choroids plexus stroma and psammoma bodies. Microsc. Res. Tech. 70, 617–627 (2007).

65. Jovanović, I., Ugrenović, S., Vasović, L., Petrović, D. & Cekić, S. Psammoma bodies - friends or foes of the aging choroid plexus. Med. Hypotheses 74, 1017–1020 (2010).

66. Živković, V.S., Stanojković, M.M. & Antić, M.M. Psammoma bodies as signs of choroid plexus ageing – a morphometric analysis. Vojnosanit. Pregl. 74, 1054–1059 (2017).

67. Montine, T.J. et al. National Institute on Aging-Alzheimer’s Association guidelines for the neuropathologic assessment of Alzheimer’s disease: a practical approach. Acta Neuropathol. 123, 1–11 (2012).

68. Schneider, C.A., Rasband, W.S. & Eliceiri, K.W. NIH Image to ImageJ: 25 years of image analysis. Nat. Methods 9, 671–675 (2012).

