## Supplementary information for "Biomarkers in cerebrospinal fluid sediments"

**Supplementary Table 1** Demographic and neuropathological data about the brain donors used for immunofluorescence. AD: Alzheimer disease; DS: own Syndrome; FTLD: frontotemporal lobar degeneration; PMD: post-mortem delay; PSP: progressive supranuclear palsy.

| Subject | Sex | Age at death | PMD (hh:mm) | Neuropathological diagnosis |
|---|---|---|---|---|
| 1 | Female | 88 | | FTLD-PSP |
| 2 | Female | 64 | | DS- AD |

**Supplementary Table 2** Primary antibodies and synthetic antigens used for immunofluorescence.

| **Primary antibodies** | | | |
|---|---|---|---|
| **Antibody** | **Clone** | **Company** | **Host** |
| anti-tau | Tau5 | Thermo Fisher Scientific; Rockford, IL, USA | Mouse |
| anti-Aβ | 12F4 | BioLegend, San Diego, California, USA | Mouse |
| anti-Aβ | 6E10 | BioLegend, San Diego, California, USA | Mouse |
| anti-p62 | 2C11 | Abcam, Cambridge, UK | Mouse |
| **Synthetic antigens** | | | |
| **Antigen** | **Company** | **Reconstitution** | |
| Aβ Protein Fragment 1-42 | Sigma-Aldrich, Merck, Darmstadt, GER | 250μg/mL in $NH_4OH$ 1% | |
| Tau-381, recombinant human | Sigma-Aldrich, Merck, Darmstadt, GER | 1 μg/μL in deionized water | |

**Supplementary Figure 1** Representative images from haematoxylin and eosin-stained sections. The first row images from an AD patient, with first two columns featuring a CA and a psammoma body in the temporal third and fourth columns showing the same features in the spinal cord. The second row shows similar imag PSP patient. Scale bar: 20 µm.

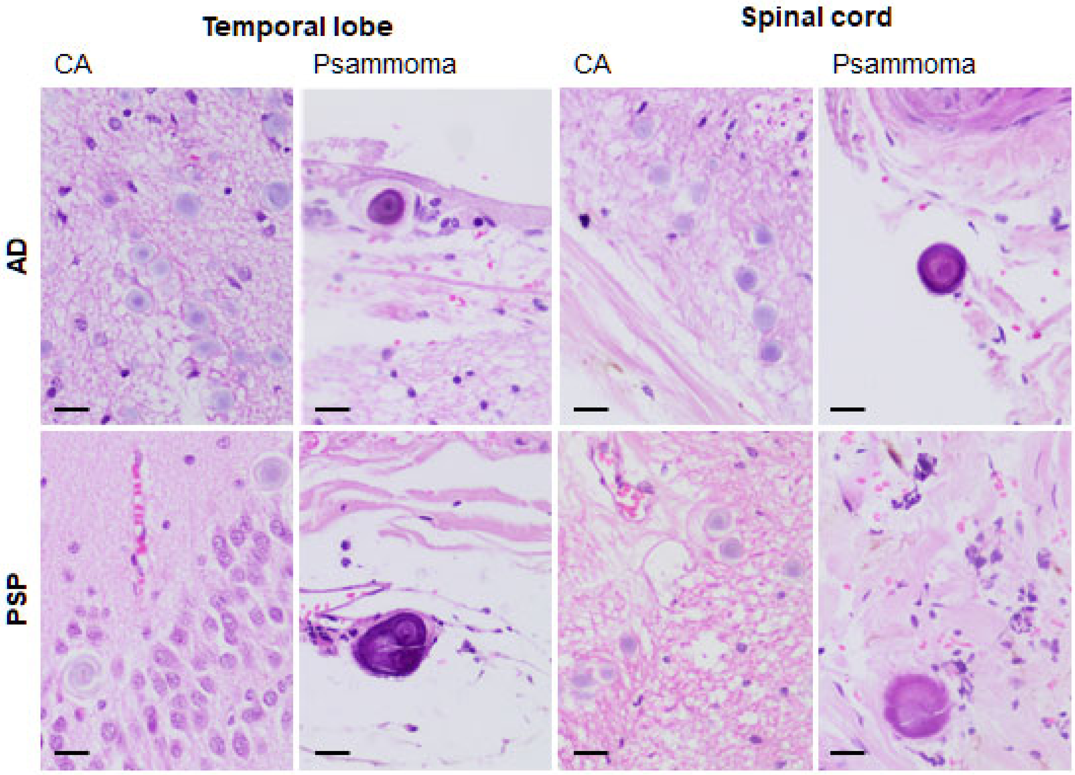


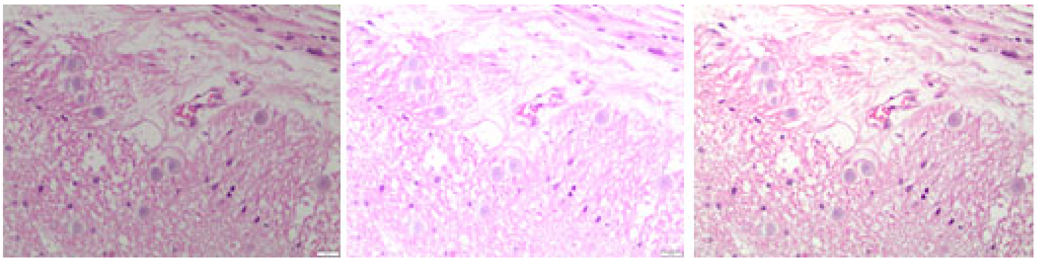
